# On multiple infections by parasites with complex life cycles

**DOI:** 10.1101/2023.10.13.562184

**Authors:** Phuong L. Nguyen, Chaitanya S. Gokhale

## Abstract

Host manipulation is a common strategy of parasites with complex life cycles. It directly affects predator-prey dynamics in trophically transmitted parasites. Theoretical studies suggest that predation-enhancing manipulation of-ten decimates the prey population, making parasites prone to extinction. Host manipulation, however, can also reduce predation due to conflicting interests when multiple parasites infect a host, which is often neglected in theoretical studies. Misaligned interests of coinfecting parasites can occur due to limited carrying capacity or parasitoid developmental stage. Including this realistic complexity in a mathematical model, the results depart from previous studies substantially. We show that coinfecting multi-trophic parasites can preserve the predator-prey system and themselves through manipulation and reproduction parameters. Our study highlights the necessity of and provides the means for incorporating the reality of multiple parasites and their multi-trophic life cycles into the theory of parasite ecology.

## Introduction

Parasites infect life on earth ubiquitously, and many of these parasites have complex life cycles (Zimmer, 2001). While a complex life cycle can be defined as abrupt ontogenic changes in morphology and ecology (Benesh, 2016), it typically involves numerous host species that a parasite needs to traverse to complete its life cycle. This complex life cycle results in the evolution of various strategies that enable successful parasite transmission from one host species to another. Host manipulation is a famous strategy that inspires many science fiction movies and novels, where a parasite can alter its host’s morphology and/or behaviour to enhance its transmission to the next host (Hughes et al., 2012). Host manipulation has been shown in many host-parasite systems, from parasites with simple life cycles to those with a complex life cycle that involves more than one host species (Hughes et al., 2012; Molyneux and Jefferies, 1986). For instance, sand flies infected by *Leishmania* parasites bite more and take more time for a blood meal from mammals (the definitive host of *Leishmania*) compared to their uninfected counterparts (Rogers and Bates, 2007). Copepods infected by cestode parasites are more active and accessible to sticklebacks (the cestodes’ definitive hosts) than uninfected copepods (Wedekind and Milinski, 1996).

Theoretical studies have long attempted to understand the ecological and evolutionary consequences of host manipulation. Roosien et al. (2013) and Hosack et al. (2008) showed that manipulative parasites could increase the disease prevalence in an epidemic. Gandon (2018) studied the evolution of the manipulative ability of infectious disease parasites, showing different evolutionary outcomes depending on whether the pathogen can control its vector or host. Hadeler and Freedman (1989); Fenton and Rands (2006) and Rogawa et al. (2018) showed that host manipulation could stabilise or destabilise the predator-prey dynamics depending on how manipulation affects the predation response function and the reproduction of the infected definitive host. Seppälä and Jokela (2008) showed that host manipulation could evolve even when it increases the risk of the intermediate host being eaten by a non-host predator, given that the initial predation risk is sufficiently low.

Most studies mentioned above have not explicitly considered a crucial aspect of parasite dynamics – multiple infections (Kalbe et al., 2002) i.e. the presence of multiple individual parasites within a single host. Multiple infections are a norm rather than an exception in parasitism. They result in the coinfection of more than one parasite inside a host, which may alter the manipulative outcomes (figure 1). An alignment of interest between coinfecting parasites may enhance manipulation, while a conflict of interest may reduce the manipulative effect. Indeed, Hafer and Milinski (2015) showed that copepods infected by two cestode parasites reduce the activity of copepods when both parasites are at the same noninfectious stage, i.e. both parasites are not ready to transmit. When two infectious parasites infect the copepods, the copepods’ activity increases, and so does the predation risk for the copepod. However, when the copepods are infected by one infectious and one noninfectious parasite, their interests clash, and the infectious parasite wins.

**Figure 1:**
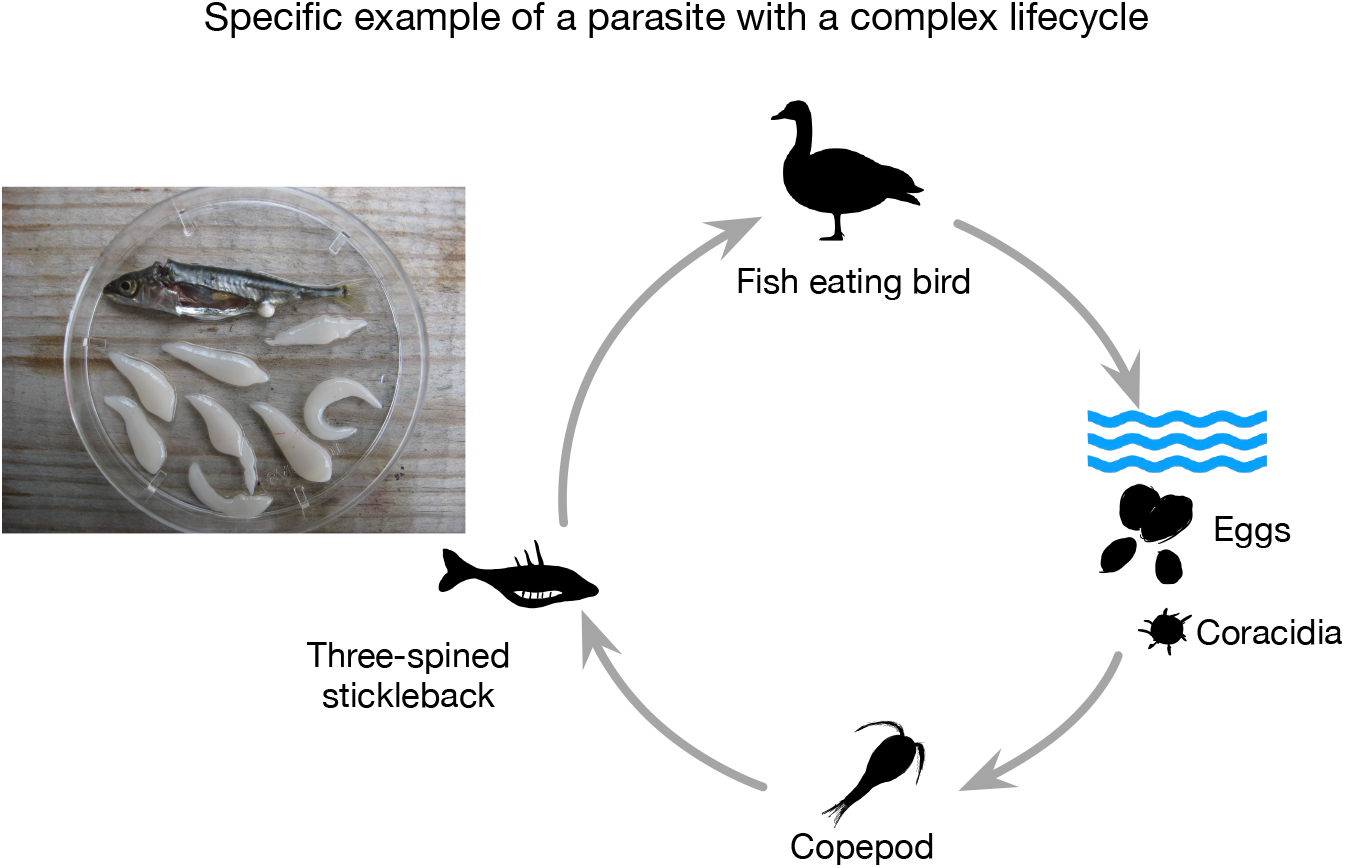
Who is in control? Schistocephalus eggs hatch into microscopically tiny swimming larvae. These larvae are eaten by copepods, where they develop to the second larval stage. However, the copepod is only the first intermediate host. The larvae are then eaten by sticklebacks, reaching the third larval stage and growing prominently in size and weight. For the parasite to successfully reach its final host, a warm-blooded animal like a bird, it manipulates its intermediate hosts. The presence of multiple parasites in the same host can lead to competition and strategic decisions about investment in manipulation and growth. Indeed, a stickleback can be infected by numerous parasites, all vying for control, as shown and photographed by Martin Kalbe (Kalbe et al., 2002). While this is a specific example of a parasite with a complex life cycle, our model abstracts the concept to generic multi-host life cycles with an environmental component.

Theoretical work that considers multiple infections often focuses on the evolution of virulence (van Baalen and Sabelis, 1995; Alizon et al., 2013; Alizon and van Baalen, 2008; Choisy and de Roode, 2010; Alizon, 2012), while host manipulation in trophically transmitted parasites receives less attention. Even though host manipulation and virulence correlate with parasite transmission, there are subtle differences, such that virulence implies an addition to the natural mortality rate of the infected host, whereas manipulation links to the immediate death of the intermediate host due to predation. Host manipulation in trophically transmitted parasites, therefore, strongly affects the entire predator-prey dynamics. Theoretical studies regarding host manipulation rarely consider multiple infections. Studies incorporating this feature neglect the predator-prey dynamics, which will likely have important feedback on the evolution of host manipulation (Parker et al., 2003; Vickery and Poulin, 2009). Moreover, these models assume that transmission from definitive hosts to intermediate hosts is due to direct contact between the two types of hosts (Rogawa et al., 2018; Hadeler and Freedman, 1989; Fenton and Rands, 2006). This is often not the case in nature, as parasites are released from the definitive hosts into the environment. Transmission thus happens only when intermediate hosts have contact with this free-living parasite pool. The inclusion of this free-living stage could have a profound effect on the dynamics of the whole predator-prey-parasite system.

Our study addresses the gap in the theoretical work on host manipulation in trophically transmitted parasites. We include multiple infections of the same parasite species and consider the dynamics of the free-living parasite pool. Our compartment model helps illustrate a parasite’s complex life cycle with two host species: an intermediate host preyed upon by a definitive host. Transmission from the intermediate host to the definitive host occurs when predation on infected intermediate hosts happens. Reproduction only happens in the definitive hosts. New parasites then enter the environment, where the cycle continues. We focus on the intermediate host manipulation, such that the parasite increases the uptake of the intermediate host by the definitive host to increase its transmission rate. We then analyse the effect of host manipulation on the ecological dynamics in the predator-prey-parasite system. We found that sabotage in host manipulation almost always pushes the dynamical system toward bistability, provided the reproduction in a single infection is sufficiently small. The bistable nature suggests that the predator-prey parasite system is finely balanced and susceptible to extinction via ecological disturbances. Initially surprising, we showed that co-operation in host manipulation and enhanced reproduction in co-infecting parasites is not always beneficial and might expose the parasite population to the risk of extinction.

## Model

Our model concerns the complex life cycle of a trophically transmitted parasite that requires two hosts: an intermediate host and a definitive host. Reproduction only happens inside the definitive hosts, releasing new parasitic progeny in the environment. An intermediate host can be infected if it encounters this free-living parasite pool. Finally, when a definitive host consumes an infected intermediate host, the definitive host gets infected, and the parasite completes its life cycle.

For simplicity, we assume that hosts can be infected by one (single infection) or, at most, two parasites of the same species (double infections). Thus, while *I*_*s*_ and *D*_*s*_ are the susceptible intermediate and definitive hosts, their singly and doubly infected counterparts are denoted by *I*_*w*_ and *D*_*w*_ and *I*_*ww*_ and *D*_*ww*_ respectively. Our model is, therefore, more relevant to the macroparasitic system. Figure (2) illustrates the transmission dynamics, and details of the model’s variables and parameters are shown in Table 1. Note that multiple infections in nature often involve more than two parasites. Typically, the number of parasites in multiple infections follows a negative binomial distribution, i.e. most hosts are infected with a few parasites, and few hosts are infected with many parasites (Wilson et al., 1996).

**Table 1:** Description of variables and parameters.

| Parameters and Variables | Description | Dimensionality |
| --- | --- | --- |
| $I_i$ | Density of intermediate hosts that are susceptible $i = s$, singly infected $i = w$, or doubly infected $i = ww$ | $[I_i]t^{-1}$ |
| $D_i$ | Density of definitive hosts that are susceptible $i = s$, singly infected $i = w$, or doubly infected $i = ww$ | $[D_i]t^{-1}$ |
| $W$ | Density of parasites released from definitive hosts into the environment | $[W]t^{-1}$ |
| $r$ | Reproduction rate of intermediate host | $t^{-1}$ |
| $k$ | Competition coefficient | $[I_i]^{-1}$ |
| $d$ | Natural death rate of intermediate hosts | $t^{-1}$ |
| $\alpha_i$ | Additional death rate of intermediate hosts due to infection by a single parasite ($i = w$) or two parasites ($i = ww$) | $t^{-1}$ |
| $p$ | Probability that two parasites co-transmit from the environment to an intermediate host | dimensionless |
| $\gamma$ | Transmission rate of parasites in the environment to intermediate hosts | $[W]^{-1}t^{-1}$ |
| $\rho$ | Baseline capture rate | $[I_i]^{-1}t^{-1}$ |
| $c$ | Coefficient of energy conversion into new definitive host | dimensionless |
| $\mu$ | Natural death rate of definitive hosts | $t^{-1}$ |
| $\sigma_i$ | Additional death rate of definitive hosts due to infection by a single parasite ($i = w$) or two parasites ($i = ww$) | $t^{-1}$ |
| $\sigma_i$ | Additional death rate of the hosts due to being infected by a single parasite ($i = w$) or two parasites ($i = ww$) | $t^{-1}$ |
| $q$ | Probability that two parasites co-transmit from intermediate hosts to definitive hosts | dimensionless |
| $\beta_i$ | Transmission rate of parasites from intermediate hosts to definitive hosts | $[I_i]^{-1}t^{-1}$ |
| $f_i$ | Reproduction rate of parasites in singly infected definitive hosts ($i = w$) or doubly infected hosts ($i = ww$) | $t^{-1}$ |
| $\delta$ | Natural death rate of parasites in the environment | $t^{-1}$ |
| $h$ | Probability that the parasites successfully established inside the definitive host | dimensionless |

\* $[I_i]$ , $[D_i]$ , and $[W]$ have the same unit (*individual area*<sup>-1</sup>)

However, since we use a compartmental model, enabling binomial distribution would mean infinitely many differential equations, making it impossible to formulate and analyze the model. Instead, we focus on another aspect of multiple infections, that is, co-transmission, which has been shown to affect the evolutionary trajectories of parasites in infectious disease (Alizon, 2012). Given an infection, the probability that two parasites from the parasite pool co-transmit to an intermediate host is denoted by *p*. Thus, 1 − *p* is the probability that a single parasite enters an intermediate host. When a definitive host consumes an intermediate host infected by two parasites, there is a probability *q* that the parasites co-transmit to the definitive host. With probability 1 − *q*, only one parasite successfully transmits. This formulation assumes that infection always happens when intermediate hosts encounter free-living parasites and when definitive hosts consume infected intermediate hosts (Figure. 2). The dynamics of a complex life cycle parasite that requires two host species is described by the following system of equations, firstly for the intermediate host as,

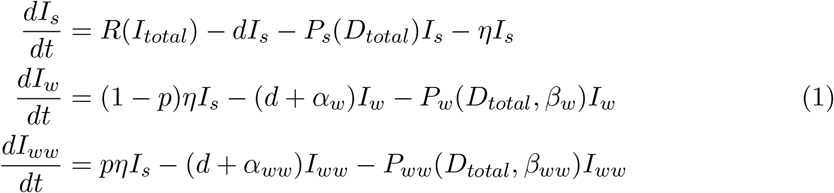

where *R*(*I*_*total*_) represents the birth rate of the intermediate hosts, a function of both infected and uninfected individuals *I*_*total*_ = *I*_*s*_ + *I*_*w*_ + *I*_*ww*_. Intermediate hosts die at a natural rate *d*, and parasites cause additional mortality rate *α*_*w*_ in single infection and *α*_*ww*_ in double infection. *P*_*s*_, *P*_*w*_, *P*_*ww*_ are the predation functions of definitive hosts on susceptible, singly infected and doubly infected intermediate hosts. The predation function depends on the density of all definitive hosts *D*_*total*_ = *D*_*s*_ + *D*_*w*_ + *D*_*ww*_ and the manipulative strategies of parasites in the intermediate hosts. In particular, if a single parasite infects an intermediate host, the manipulation strategy is *β*_*w*_. However, if the intermediate host is co-infected, the manipulation strategy is *β*_*ww*_. We assume no specific relationship between *β*_*w*_ and *β*_*ww*_ to explore all possible ecological outcomes of the system. The force of infection by parasites in the environment is denoted by *η* = *γW*, where *γ* represents the infection rate of free-living parasites. The force of infection is a term often used in epidemiology, which represents the rate at which a host gets infected by the parasites. Since parasites can manipulate intermediate and definitive hosts, whenever we mention host manipulation, it specifically refers to the manipulation in intermediate hosts, which correlates to the predation rate.

**Figure 2:**
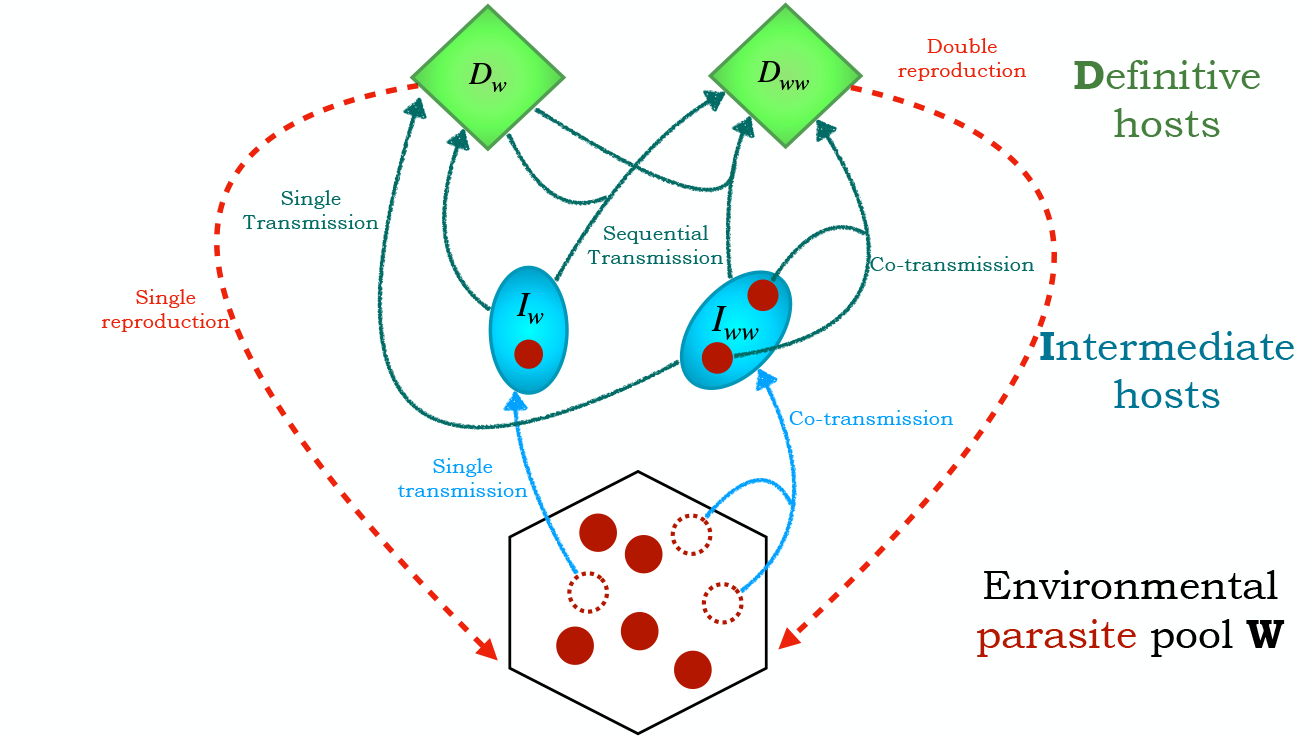
Schematics of the transmission routes. Blue ovals represent the intermediate hosts, while green diamonds represent the definitive hosts. The hexagon represents the parasite pool compartment, with the red circles illustrating the free-living individual parasites. The parasites infect the intermediate hosts singly (*I*_*w*_) or doubly (*I*_*ww*_) upon encounter between the intermediate hosts and the parasite pool (blue arrows). These intermediate hosts are then predated upon by the definitive hosts (green arrows), thus moving the parasites to the final host (either as *D*_*w*_ or *D*_*ww*_) where they can reproduce and reenter the free-living stage in the environmental pool *W* (red dashed arrows).

For the definitive hosts, we have,

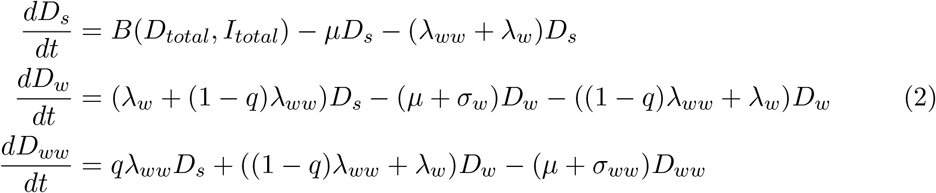

where *B*(*D*_*total*_, *I*_*total*_) represents the birth rate of definitive hosts. The birth rates depend on the density of both intermediate and definitive hosts, infected or uninfected. The natural mortality rate of definitive hosts is represented by *µ*, and parasites induce additional mortality rates *σ*_*w*_ and *σ*_*ww*_ in single and double infection, respectively. The force of infection that corresponds respectively to singly infected intermediate host (*I*_*w*_) and doubly infected intermediate hosts (*I*_*ww*_) is denoted respectively by *λ*_*w*_ = *h*(*ρ*+*β*_*w*_)*I*_*w*_ and *λ*_*ww*_ = *h*(*ρ*+*β*_*ww*_)*I*_*ww*_, where *ρ* is the baseline predation rate, i.e. the basic constitutive level of predation, and *h* is the probability that the parasite successfully establishes inside the host. Without manipulation, that is, *β*_*w*_ = *β*_*ww*_ = 0, the parasite is still transmitted via the baseline predation *ρ*. The dynamics of the free-living parasites in the environment are then given by,

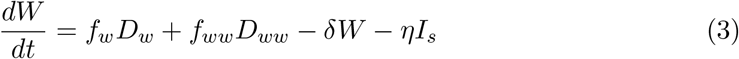

where *f*_*w*_ and *f*_*ww*_ are the reproduction rates of parasites in single and double infection, respectively, and parasites die naturally at a rate *δ*.

Here, we focus on manipulation that enhances transmission from intermediate hosts to definitive hosts; we thus simplify the transmission from the parasite pool to intermediate hosts so that no sequential infection occurs. This assumption is motivated because the prey’s life cycle is often shorter than the predator’s. A prey likely encounters the free-living parasite pool once and then dies due to predation, making sequential transmission less likely at this state. Sequential infection can happen when parasites transmit from intermediate hosts to definitive hosts. Therefore, a singly infected definitive host can be further infected by another parasite if it consumes infected intermediate hosts.

### Basic reproduction ratio *R*_0_ of the parasites

The basic reproduction ratio *R*_0_ (or basic reproduction number as often used in epidemiology) indicates parasite fitness. It can be understood as the expected number of offspring a parasite produces during its lifetime when introduced to a susceptible host population. We calculate the basic reproduction ratio *R*_0_ using the next-generation method (Diekmann et al., 1990, 2009; Hurford et al., 2010) (See SI1 for details).

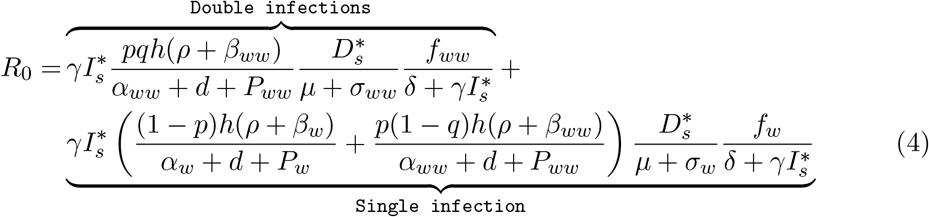

where 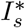 and 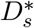 are the densities of susceptible intermediate and definitive hosts at the disease-free equilibrium. Here, the expression of *R*_0_ contains the possible reproduction routes of a parasite, which can be via double or single infections. The first component corresponds to the double infections route, in which the focal parasite co-transmits with another parasite into a susceptible intermediate host, then co-transmits into a susceptible definitive host and reproduces. Here, parasites are so rare that only co-transmission matters and the compartments with sequential infections are neglected. The second component corresponds to the single infection route, wherein the focal parasite infects a susceptible intermediate host via single or double infections. The parasite then transmits alone into the susceptible definitive host and eventually reproduces.

If *R*_0_ *>* 1, a parasite spreads when introduced into the disease-free equilibrium of prey and predator. Intuitively, the higher the density of susceptible intermediate and definitive hosts, the larger the value of *R*_0_ as the infection reservoir is more extensive. In contrast, regardless of the explicit form of the predation function, the higher the predation rate *P*_*w*_ and *P*_*ww*_, the lower the value of *R*_0_ given the smaller reservoir of intermediate hosts. The effect of host manipulation on the value of *R*_0_ is more complex; as host manipulation becomes efficient, the transmission rate from the intermediate host to the definitive host increases, but so does the predation rate. A higher predation rate results in a smaller intermediate host reservoir for the parasites to infect. To understand the effect of manipulation on parasites’ fitness and the system’s ecological dynamics, we next specify the predation functions. We consider linear functions for predation to begin with,

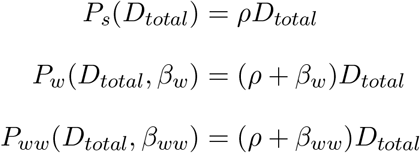

where *ρ* is the baseline capture rate of the predator on the prey. If an intermediate host is infected, it is captured by the definitive hosts with rate *ρ* + *β*_*w*_ if it is singly infected and with rate *ρ* + *β*_*ww*_ if it is doubly infected. Zero values for *β*_*w*_ and *β*_*ww*_ suggest no manipulation, and predation is at the baseline value *ρ*.

For simplicity, we also consider a linear function of the birth of definitive hosts

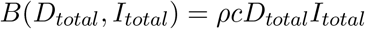

where *c* is the efficiency of converting prey into predator’s offspring. It is important to note that host manipulation affects population dynamics via its influence on the predation rate, not the physiological aspect of the definitive host, i.e., the predator. The birth rate of the predators thus depends on the capture rate, but it is not affected by host manipulation; to our best knowledge, there is no supporting evidence to consider otherwise.

The explicit form of 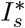 and 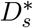, capturing the predator-prey dynamics, depends on the precise form of all birth and predation functions *B, R, P*_*s*_, *P*_*w*_ and *P*_*ww*_. However, it does not depend on the ability to manipulate or any other parameter of the parasite. Given that the birth rate of the predator and the predation rate are linear functions in prey and predator density, the form of the birth rate *R* of the prey has a significant effect on the susceptible intermediate and definitive host dynamics.

### Birth function of intermediate hosts

The simplest form of the prey’s birth rate is a linear function, in which case the disease-free equilibrium is always in a cyclic regime (see SI 2). This follows from the Lotka-Volterra system using linear functions for prey birth and predation (Lotka, 1920). Since the disease-free dynamics is cyclic, it is difficult to analyse the spread of a parasite using the basic reproduction ratio, which is evaluated when the disease-free state is stable. Here, *R*_0_ *>* 1 happens when *γ*, the transmission rate from the environment to intermediate hosts, and the reproduction rates *f*_*w*_, *f*_*ww*_ are quite large (as compared to the theoretical threshold shown by the mathematical conditions in SI 3). However, even when this condition is satisfied, the parasite may not be able to spread and persist in cyclic susceptible host dynamics (Figure SI.1). This result agrees with the conclusion in (Ripa and Dieckmann, 2013), which suggests that it is difficult for a mutant to invade a cyclic resident population. In our case, it is not the invasion of a mutant in a resident population but the invasion of a parasite in a cyclic disease-free host population; the argument, however, remains valid in both cases. This issue deserves a more thorough investigation, which is out of the scope of this article. Therefore, we choose a non-linear birth function of the intermediate hosts to obtain a stable disease-free state and focus on the effect of host manipulation on the ecological dynamics (Figure 3).

**Figure 3:**
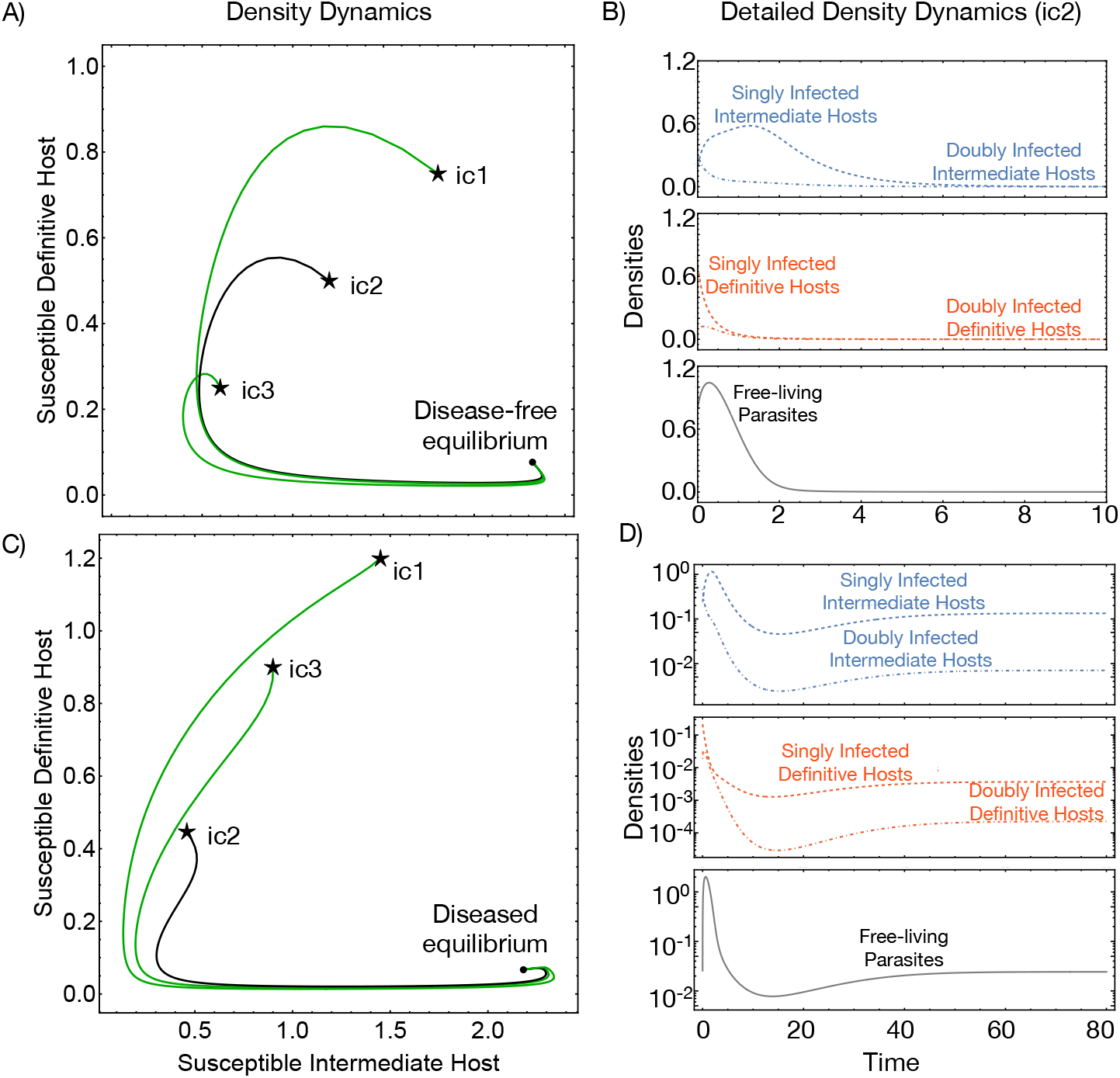
Ecological dynamics of the predator-prey-parasite system. On the left, we show the density dynamics of the susceptible intermediate and definitive hosts at different initial conditions (ic1, ic2, and ic3). The detailed dynamics of infected compartments are further shown for specific initial conditions (ic2), including the free-living parasite dynamics. A-B) A case of a disease-free equilibrium being reached from different initial conditions (ic). C-D) A case where the parasite survives. Parameters for disease free equilibrium *ρ* = 1.2, *d* = 0.9, *r* = 2.5, *γ* = 2.9, *α*_*w*_ = *α*_*ww*_ = 0, *β*_*w*_ = *β*_*ww*_ = 1.5, *p* = 0.05, *c* = 1.4, *µ* = 3.9, *σ*_*w*_ = *σ*_*ww*_ = 0, *q* = 0.05, *f*_*w*_ = *f*_*ww*_ = 7.5, *δ* = 0.9, *k* = 0.26, *h* = 0.6. Disease stable equilibrium has the same parameter values except for higher host manipulation *β*_*w*_ = *β*_*ww*_ = 4.5 and parasite reproduction *f*_*w*_ = *f*_*ww*_ = 45

The logistic growth for the non-linear birth function follows by

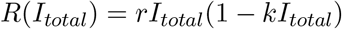

where *r* is the intrinsic growth rate of the intermediate hosts, and *k* is the intraspecific competition coefficient. The disease-free equilibrium is as follows,

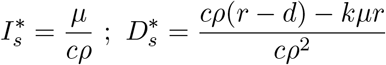

This equilibrium is positive and stable if components of the parasite, such as reproduction and transmission, are sufficiently small; details of the condition can be found in section SI 4. Here, as the reproduction and transmission values of the parasite are not sufficient, it goes extinct (Figure 3A, B), leaving the predator-prey dynamics attaining equilibrium (Figure 3C, D)

When a parasite appears in the disease-free equilibrium, it spreads if its reproduction ratio *R*_0_ *>* 1 (Figure 4). Since the expression is complicated, we could only obtain analytical solutions for this inequality with assumptions. We assume the same parasite virulence, *α*_*w*_ = *α*_*ww*_, *σ*_*w*_ = *σ*_*ww*_, and reproduction in double infection as a linear function concerning reproduction in single infections, *f*_*ww*_ = *ϵf*_*w*_. When *ϵ >* 1, reproduction in double infections is enhanced compared to in single infections, whereas for *ϵ* ≤ 1, it is suppressed or equal to reproduction in single infections. We found that the parasite can establish if its reproduction value in a single infection *f*_*w*_ is more significant than a threshold (Figure 5, see section SI 5 and Eq. (SI.19)).

**Figure 4:**
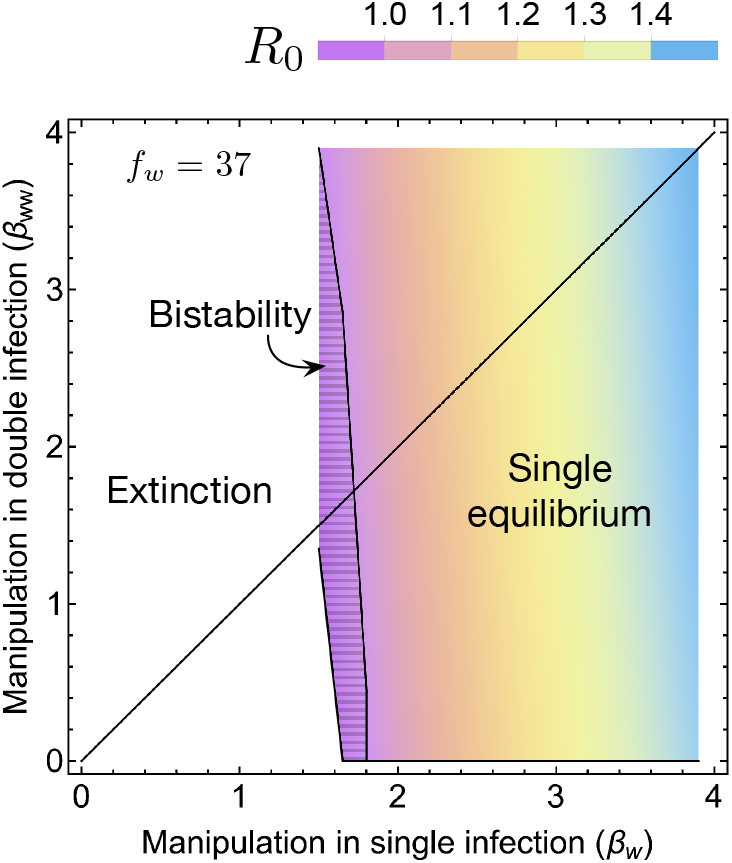
Effect of manipulation in single and double infections on the reproduction ratio. *R*_0_. *R*_0_ values increase with more efficient manipulation in single and double infection. The hatched area indicates the bistable region. As manipulation in single infection increases, the system only has one stable equilibrium. On the black line, the manipulation level is equal between single and double infection (*β*_*w*_ = *β*_*ww*_). In the upper triangular area, parasites cooperate, and in the lower triangular area, parasites sabotage. Common parameter: *ρ* = 1.2, *d* = 0.9, *r* = 2.5, *γ* = 2.9, *α*_*w*_ = 0, *α*_*ww*_ = 0, *p* = 0.05, *c* = 1.4, *µ* = 3.9, *σ*_*w*_ = 0, *σ*_*ww*_ = 0, *q* = 0.05, *δ* = 0.9, *k* = 0.26, *f*_*w*_ = 37, *ϵ* = 4.5, *h* = 0.6.

**Figure 5:**
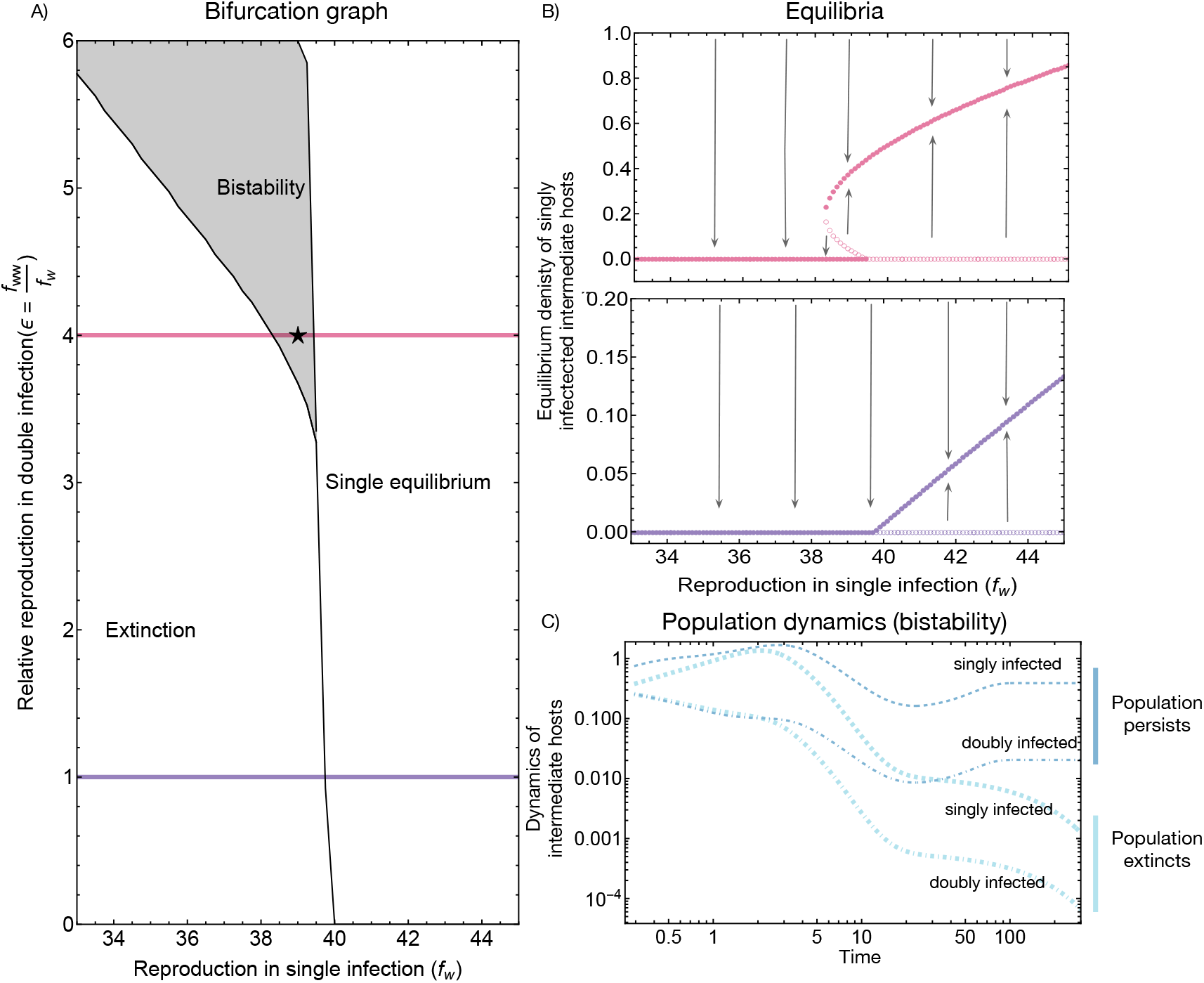
Effect of parasite reproduction on the ecological dynamics. A) A bifurcation graph for different reproduction values in single and double infections. B) Equilibrium density of intermediate host when *ϵ* = 4 when bistability occurs at high values of *f*_*w*_ (in pink), and *ϵ* = 4 when only one stable equilibrium exists at high values of *f*_*w*_ (in purple). C) Details of the parasite population dynamics in the case of bistability shown through the infected intermediate hosts. When the parasites start at high density, the parasite population persists, whereas when they start at lower density, they perish. Filled circles indicate stable equilibria, and open circles indicate unstable equilibrium. Parameter *ρ* = 1.2, *d* = 0.9, *r* = 2.5, *γ* = 2.9, *α*_*w*_ = 0, *α*_*ww*_ = 0, *β*_*w*_ = 1.5, *β*_*ww*_ = 1.5, *p* = 0.05, *c* = 1.4, *µ* = 3.9, *σ*_*w*_ = 0, *σ*_*ww*_ = 0, *q* = 0.05, *δ* = 0.9, *k* = 0.26, *h* = 0.6.

Our numerical results show that the parasite reproduction is substantial compared to other parameters (Figure 5A). For instance, in the parameter set used to generate Figure 5, to spread in the predator-prey system, the value of parasite reproduction (*f*_*w*_) has to be at least 20 times the value of intermediate host reproduction *r* = 2.5, given that both these parameters represent the *per capita* growth rate of the parasite and the intermediate host population. This observation suggests that trophically transmitted parasites should release many offspring into the environment to persist. Interestingly, bistability occurs if the reproduction rate of the parasite in double infections is enhanced. Bistability suggests that the parasite population is vulnerable to extinction. Specifically, if sufficient parasites are introduced into the disease-free predator-prey populations, the parasite population persists and reaches a stable equilibrium. In contrast, if only a few parasites are introduced into the disease-free populations, or if sufficient disturbance occurs when the parasite population is already established, the parasite population could go extinct (Figure 5C).

### The effect of host manipulation on ecological dynamics

Host manipulation can be cooperative; two parasites increase the predation rate on intermediate hosts, or *β*_*ww*_ *> β*_*w*_ (Hafer and Milinski, 2015). However, it can also be uncooperative; the predation rate on doubly-infected intermediate hosts is lower than that on singly-infected ones or *β*_*ww*_ *< β*_*w*_ (Hafer and Milinski, 2015). Cooperation in parasite manipulation increases the parasite’s basic reproduction ratio *R*_0_, but the manipulation in a single infection substantially affects the value of *R*_0_ (Figure 4, see section SI 6 for analytical results). Intuitively, if the manipulation in a single infection is minor, there is not enough transmission, and the parasite goes extinct. However, we could suppose that the ability to manipulate the host in a single infection is enough for the parasite population to escape extinction. In that case, the system is in a bistable state where intermediate cooperation in host manipulation cannot guarantee a single equilibrium (Hatched area Figure 4). In the bistable region, the basic reproduction ratio can be less than one, implying that the parasite with manipulative values within this range, i.e. weak manipulation ability, cannot spread. This is due to the Allee effect (Stephens et al., 1999) where the parasite spreads and persists if, initially, there are sufficient parasites in the free-living pool as well as in the intermediate and definitive hosts. On the contrary, if the initial populations of parasites are insufficient, the parasite will perish (Figure SI.2). In addition, when the system encounters bistability, the parasite population risks extinction if there is a disturbance in the community. In the following parts, we will explore scenarios where bistability may occur.

Besides manipulation, co-infecting parasites can influence each other in different life history traits. Parasites can have an enhanced reproduction rate in coinfections, i.e. *f*_*ww*_ *> f*_*w*_ (upper part of the horizontal line in all panels Figure 6). Likewise, they can compete for resources, so reproduction in double infection is suppressed compared to single infection (lower parts of the horizontal lines in all panels Figure 6). Without any assumption on the link between manipulative ability and reproduction, and a linear relationship between manipulation in single and double infections, we explore all possible combinations of cooperation-sabotage range in manipulation and suppressed-enhanced range in reproduction. This results in four scenarios of parameter combinations: i, parasites sabotage manipulation but have enhanced reproduction – manipulative incoordination (top left quadrants in all panels Figure 6), ii, parasites cooperate to increase manipulation and enhance reproduction – coordination (top right quadrants in all panels Figure 6), iii, parasites cooperate in manipulation but suppress reproduction – reproductive incoordination (bottom right quadrants in all panels Figure 6), and iv, parasites sabotage manipulation and suppress reproduction – discordance (bottom left quadrants in all panels Figure 6).

**Figure 6:**
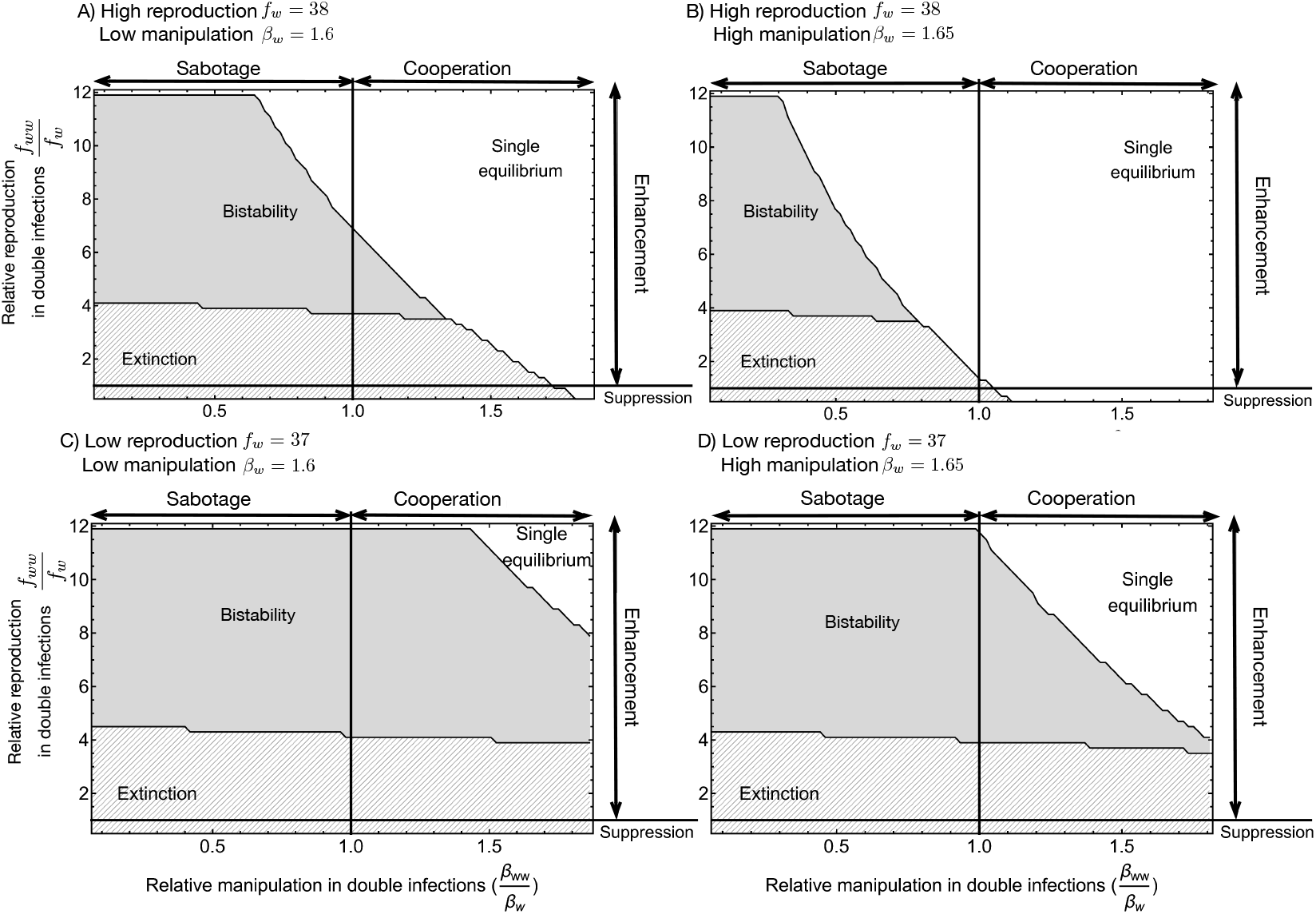
Effect of manipulation and reproduction on bistability. The bistability area (shaded areas) reduces as the reproduction rate (*f*_*w*_) and manipulation (*β*_*w*_) in a single infection increases. Reproduction in single infection decreases from the upper panels (A, B) to the lower panels (C, D), while manipulation in single infection increases from the left panels (A, C) to the right panels (B, D). Manipulation and reproduction levels are equal between single and double infection on the vertical and horizontal lines. On the left side of the vertical line, *β*_*ww*_ *> β*_*w*_, indicating cooperation, whereas on the right side of the vertical line, *β*_*ww*_ *< β*_*w*_, indicating sabotage. On the upper part of the horizontal line, *f*_*ww*_ *> f*_*w*_, indicating enhanced reproduction, whereas, on the lower part of the horizontal line, *f*_*ww*_ *< f*_*w*_, indicating suppressed reproduction. Common parameter: *ρ* = 1.2, *d* = 0.9, *r* = 2.5, *γ* = 2.9, *α*_*w*_ = 0, *α*_*ww*_ = 0, *p* = 0.05, *c* = 1.4, *µ* = 3.9, *σ*_*w*_ = 0, *σ*_*ww*_ = 0, *q* = 0.05, *δ* = 0.9, *k* = 0.26, *β*_*w*_ = 1.65, *h* = 0.6.

If coinfected parasites are discordant, i.e. uncooperative in manipulations and show suppressed reproduction, they cannot persist (bottom left quadrants Figure 6A-D). On the other extreme, where they are highly cooperative in manipulation and show enhanced reproduction, i.e., an extreme level of coordination, there is a guaranteed single equilibrium for parasite existence (top right quadrants Figure 6A-D). Note that this happens at the combination of *β*_*ww*_*/β*_*w*_ → ∞ and *f*_*ww*_*/f*_*w*_ → ∞, a scenario that is rather impossible in reality. We often expect intermediate levels of coordination where a bistable area could occur (top right quadrant in Figure 6A, C, D). However, the size of this area is sensitive to the value of reproduction and manipulation in a single infection. In particular, higher values of these two parameters reduce the bistability area so that sufficiently large reproduction in a single infection can guarantee single equilibrium when parasites coordinate (Figure 6 B, D). In contrast, slightly reducing values of either reproduction or manipulation in a single infection increases the bistability area (Figure 6A, C, D). If the parasites sabotage each other, the system is highly prone to bistability and only has a single equilibrium when reproduction is enhanced. Interestingly, reproductive incoordination, with depressed reproduction and sufficient manipulative cooperation, always leads to a single equilibrium of the system (bottom right quadrants Figure 6, note that if we extend the relative manipulation in Figure C and D, we also obtain single equilibrium in this area). While a single equilibrium guarantees the existence of a parasite population, bistability indicates that a disturbance of the system may likely lead to the extinction of the parasite population. This suggests that the benefits of coordination in reproduction and manipulation are context-dependent. Coordinating is advantageous if no significant tradeoffs and reproduction or manipulation in single infections are large enough.

We now explore the effect of co-transmission probability on the bistability of the system (Figure 7). First, extinction is more likely with varying levels of co-transmission from the parasite pool to the intermediate host, *p*, compared to varying levels of co-transmission from the intermediate host to the definitive host, *q*. For exceptionally high levels of cooperation and intermediate values of *p* and *q*, the predator-prey-parasite system will always persist with one stable equilibrium. However, limitations and trade-offs are often unavoidable, and such high values of cooperation may be impossible, putting the system in the parameter space where bistability likely occurs. When the parasite sabotages manipulation, the bistable area decreases with increasing *p* and *q*. However, this bistable area disappears with high values of *q* but not with high values of *p*. When parasites cooperate in manipulation, reducing *p* almost always leads to bistability, whereas reducing *q* can lead to a single equilibrium if cooperation is sufficiently large. Bistability indicates vulnerability to disturbance, so cooperation in manipulation may be beneficial when *q*, the co-transmission from the intermediate host to the definitive host, decreases. However, cooperation in manipulation may still harm the population by reducing *p*, the co-transmission from the parasite pool to the intermediate host.

**Figure 7:**
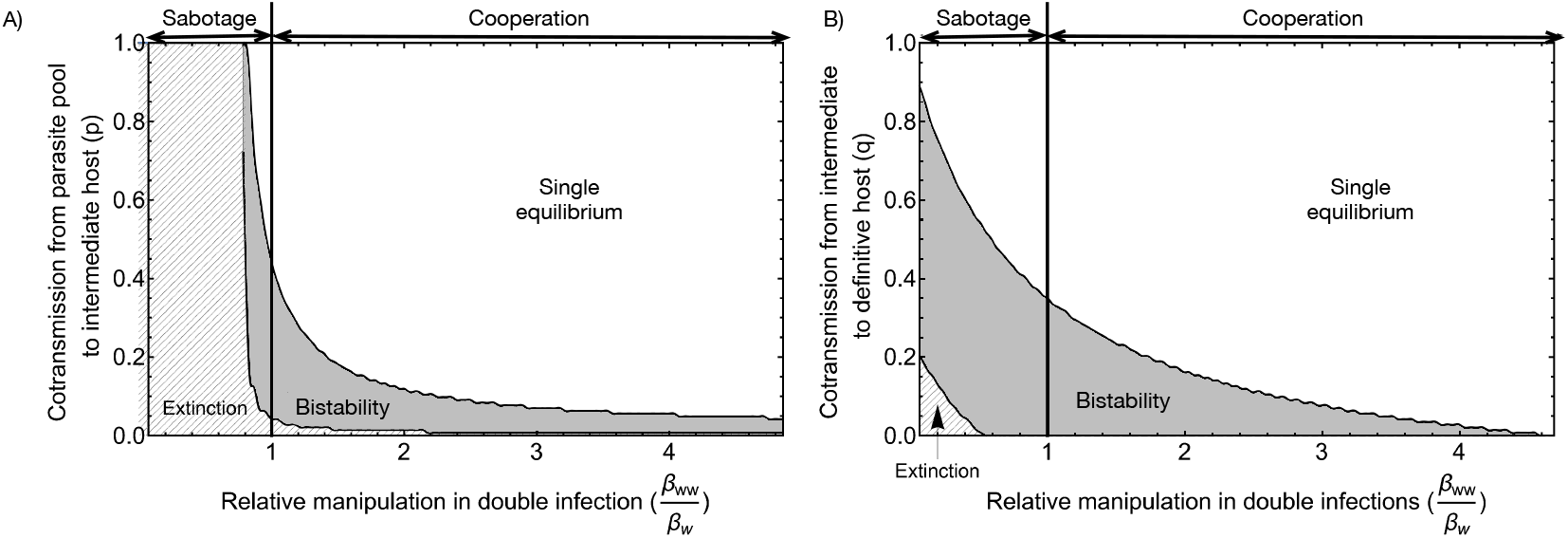
A) Effect of cotransmission from parasite pool to intermediate host. B)Effect of co-transmission from intermediate to the definitive host. On the left side of the vertical line, *β*_*ww*_ *> β*_*w*_, indicating cooperation, whereas on the right side of the vertical line, *β*_*ww*_ *< β*_*w*_, indicating sabotage. Common parameters: *ρ* = 1.2, *d* = 0.9, *r* = 2.5, *γ* = 2.9, *α*_*w*_ = 0, *α*_*ww*_ = 0, *p* = 0.05, *c* = 1.4, *µ* = 3.9, *σ*_*w*_ = 0, *σ*_*ww*_ = 0, *q* = 0.05, *δ* = 0.9, *k* = 0.26, *ϵ* = 4.5, *β*_*w*_ = 1.45, *f*_*w*_ = 38, *h* = 0.6.

## Discussion & Conclusion

Host manipulation is a ubiquitous phenomenon suggested to affect predator-prey dynamics in trophically transmitted parasites. In particular, manipulation of infected intermediate hosts to increase the predation rate of definitive hosts may result in a heavy burden of predators on the intermediate host population. This pressure can make parasites more vulnerable to extinction (Hadeler and Freedman, 1989; Fenton and Rands, 2006).

Our model shows that parasites cannot spread quickly in a cyclic predator-prey system. This delay is an expected result since even though the parasite’s basic reproduction ratio *R*_0_ is larger than one, it is estimated at the predator and prey’s unstable equilibrium (or cyclic equilibrium). Thus, when the density of the prey and predator is at the minimum value of the cycle, the “effective” *R*_0_ of the parasite can be smaller than one. Another interesting result is that the reproduction value is much larger than other parameter values, such as the *per capita* reproduction rate of the intermediate host. This result is likely due to the introduction of a free-living parasitic pool. Our model shows that in making the system more realistic, we also obtain a more realistic quantitative value for parasitic reproduction.

In the study by Rogawa et al. (2018), a non-manipulative parasite can invade a susceptible prey-predator population and cause the system to cycle. The system stops cycling and approaches a fixed point when the parasite becomes manipulative, and this stability increases with increased manipulation. In our model, non-manipulative parasites cannot persist in the system, and the parasite never leads to cyclic dynamics. These results may contradict with Rogawa et al. (2018), where non-manipulative parasites can still exist via cyclic behaviour. We suggest that the different results may be due to our introduction of a parasite pool and multiple infections, unlike the model of Rogawa et al. (2018). In their system, transmission from the definitive host to the intermediate host was assumed to result from direct contact between the two host species. Such immediate transmission could directly accelerate the feedback loop between prey and predator. Hence, faster predator-prey dynamics occur, which may lead to cyclic dynamics when parasites are introduced.

Another study on host manipulation, Iritani and Sato (2018), showed that manipulative parasites persist if they switch from suppressing to boosting predation rate. This theoretical work modelled the ability to change the manipulative strategy of a single parasite inside a host, which can be equal to introducing the developmental state of a parasite, where a suppressed predation rate protects the parasites that are not ready to transmit. That is why decreasing manipulative ability is beneficial and prevents parasite extinction. In our model, sabotaging manipulation also reduces manipulative ability, which only reduces the basic reproduction ratio *R*_0_ and makes the system bistable, exposing the parasite to the risk of extinction. This result contrasts with Iritani and Sato (2018) because in our model, the parasite cannot switch its manipulative strategy, and sabotage decreases the transmission rate from intermediate to definitive host and does not benefit the parasite in any way.

In our study, population dynamics exhibit bistability under certain circumstances. This is very likely due to the introduction of co-transmission, which has been shown to result in bistable population dynamics in plant virus Allen et al. (2019) and infectious diseases Gao et al. (2016). In this bistability region, if the system is disturbed (e.g. migration of the intermediate or definitive hosts or predation of intermediate hosts by other predators), then the density of the infected hosts may crash, leading to parasite extinction. The bistability region widens as parasites show enhanced reproduction but sabotage manipulation. This extension is because the density of the doubly infected hosts is always much smaller than the singly infected hosts, limited by sequential transmission and a small probability of cotransmission. If manipulation in a single infection is insufficient, then the transmission of the parasites depends mainly on the doubly infected hosts, which is rare. So, extinction is possible if manipulation in double infections is low.

Our study specifically focuses on the ecological dynamics of a trophically transmitted parasite between two host species. In nature, parasites with complex life cycles can have more than two hosts. However, our model of a single intermediate host species already includes enough complexity to discuss the relationship between transmission and manipulation. In addition, we consider multiple infections of the same parasite species. Although, in nature, a host can be co-infected by multiple parasites of the same or different species, the results of our model stay valid as the key aspect in host manipulation is the alignment or conflict of interest between co-infecting parasites. Here, we introduce more realistic features compared to previous models, such as a free-living parasite pool and multiple infections, regardless of some simplifications, such as multiple infections being limited to at most two parasites. Thus, we can obtain analytical results of the reproduction ratio and mathematical expressions for the existing condition of the parasite.

Our model serves as a groundwork for future exploration into more complex and realistic systems, where numerical simulation may be the only possible approach. Given that few studies considered measuring different parameters of trophically transmitted parasites (Seppälä et al., 2004; Gopko et al., 2015), our model calls for additional empirical work to measure relevant parameters, especially those from the parasite’s perspective. For instance, comparing the reproduction rate of parasites in single versus multiple infections (parameters *f*_*w*_ and *f*_*ww*_) sheds light on parasite cooperation in definitive hosts. Studying the distribution of parasites in the environment (the variable *W*) informs us about feeding strategies and reflects the distribution of parasites within intermediate hosts. Finally, comparing host condition (parameters *α*_*w*_, *α*_*ww*_, *σ*_*w*_, and *σ*_*ww*_) between no infection, single and multiple infections illustrates the magnitude of parasite virulence. Although parasite virulence has been quantified in some studies, none have examined differences between single and multiple infections. Eventually, the results of our ecological model are a baseline for further investigation of the evolution of host manipulation, where introducing the parasite pool may create interesting eco-evolutionary feedback to the system.

## Acknowledgements

We thank the Swiss National Science Foundation Sinergia grant no CRSII5 202290 for supporting Phuong Nguyen to work on this manuscript as a side project. Funding from Bayerische Forschungsallianz and from the Max Planck Society is gratefully acknowledged (CSG). We also thank Tina Scheibe for helpful discussions. This paper is dedicated to the memory of Martin Kalbe, an exceptional parasitologist who made remarkable contributions to the field. Beyond his academic prowess, Martin was a truly exemplary friend and human being, touching the lives of those around him with his warmth, kindness, and unwavering support.

