## Supplementary for "On multiple infections by parasites with complex life cycles"

### SI 1. Reproduction ratio $R_0$

The reproduction ratio of the parasite is derived from the dynamical system of the parasite, which only includes infected intermediate and definitive hosts and the free-living parasite pool. The dynamical system can be written in matrix form as follows:

$$\frac{d\mathbf{n}}{dt} = \mathbf{M}\mathbf{n}$$

where $\mathbf{n}$ is the vector of singly and doubly infected intermediate hosts, singly and doubly infected definitive hosts and free-living parasites $(dI_w, I_{ww}, D_w, D_{ww}, W)$ and $\mathbf{M}$ is the matrix that describes the dynamics

$$\mathbf{M} = \begin{pmatrix} -d-\alpha_w-P_w & 0 & 0 & 0 & (1-p)\gamma I_s \\ 0 & -d-\alpha_{ww}-P_{ww} & 0 & 0 & p\gamma I_s \\ h(\beta_w+\rho)D_s & h(\beta_{ww}+\rho)(1-q)D_s & -\lambda_w-(1-q)\lambda_{ww}-\mu-\sigma_w & 0 & 0 \\ 0 & h(\beta_{ww}+\rho)qD_s & \lambda_w+(1-q)\lambda_{ww} & -\mu-\sigma_{ww} & 0 \\ 0 & 0 & f_w & f_{ww} & -\delta-\gamma I_s \end{pmatrix}$$

The matrix $\mathbf{M}$ can be written as $\mathbf{M} = \mathbf{F} - \mathbf{V}$, where

$$\mathbf{F} = \begin{pmatrix} 0 & 0 & 0 & 0 & 0 \\ 0 & 0 & 0 & 0 & 0 \\ 0 & 0 & 0 & 0 & 0 \\ 0 & 0 & 0 & 0 & 0 \\ 0 & 0 & f_w & f_{ww} & 0 \end{pmatrix}$$

is the matrix in which its elements are the reproduction contribution of one compartment to the other compartments in the next generation, and

$$\mathbf{V} = \begin{pmatrix} \alpha_w + d + P_w & 0 & 0 & 0 & -(1-p)\gamma I_s \\ 0 & \alpha_{ww} + d + P_{ww} & 0 & 0 & -p\gamma I_s \\ -h(\rho+\beta_w)D_s & -h(\rho+\beta_{ww})(1-q)D_s & \lambda_w + \lambda_{ww}(1-q) + \mu + \sigma_w & 0 & 0 \\ 0 & -h(\rho+\beta_{ww})qD_s & -\lambda_w - \lambda_{ww}(1-q) & \mu + \sigma_{ww} & 0 \\ 0 & 0 & 0 & 0 & \delta + \gamma I_s \end{pmatrix}$$

is the matrix in which its elements include death rates or transition rates from one compartment to the others (Diekmann et al., 1990, 2009; Hurford et al., 2010).

The reproduction ratio $R_0$ is then the leading eigenvalue of the matrix $\mathbf{F}.\mathbf{V}^-1$, evaluated at the disease-free equilibrium of the intermediate and definitive hosts $I_s^*$, $D_s^*$, and $I_w = I_{ww} = D_w = D_{ww} = 0$.

### SI 2. Equilibrium stability - linear birth function for intermediate hosts

The Jacobian matrix of the system of equations (1), (2), and (3), as given in the main text, is evaluated at the disease-free equilibrium, and $B(D_s, D_w, D_{ww}, I_s, I_w, I_{ww}) = \rho c D_{total} I_{total}$ is

$$
\begin{pmatrix}
0 & r & r & -\frac{\mu}{c} & -\frac{\mu}{c} & -\frac{\mu}{c} & -\frac{\gamma\mu}{c\rho} \\
0 & -\alpha_w + \frac{(\beta_w+\rho)(d-r)}{\rho} - d & 0 & 0 & 0 & 0 & \frac{\gamma\mu(1-p)}{c\rho} \\
0 & 0 & -\alpha_{ww} + \frac{(\beta_{ww}+\rho)(d-r)}{\rho} - d & 0 & 0 & 0 & \frac{\gamma\mu p}{c\rho} \\
-c(d-r) & \frac{\beta_w(d-r)}{\rho} - c(d-r) & \frac{\beta_{ww}(d-r)}{\rho} - c(d-r) & 0 & \mu & \mu & 0 \\
0 & -\frac{\beta_w(d-r)}{\rho} & -\frac{\beta_{ww}(1-q)(d-r)}{\rho} & 0 & -\mu-\sigma_w & 0 & 0 \\
0 & 0 & -\frac{\beta_{ww}q(d-r)}{\rho} & 0 & 0 & -\mu-\sigma_{ww} & 0 \\
0 & 0 & 0 & 0 & f_w & f_{ww} & -\frac{\gamma\mu}{c\rho} - \delta
\end{pmatrix}
$$

This jacobian has seven eigenvalues, two of which have explicit expressions as $\pm\sqrt{d-r}$. Here, we always have $r > d$ to keep the equilibrium positive. Therefore, these two eigenvalues are always pure imaginary. We cannot obtain the explicit expression of the other five eigenvalues, but the dynamics remain unstable regardless of their values.

### SI 3. Invasion of parasite - Linear birth function

$R_0 > 1$ when the transmission rate from the parasite pool to intermediate hosts satisfies

$$\gamma > \frac{c\delta\rho(\mu+\sigma_w)(\mu+\sigma_{ww})(\beta_w(r-d)+\rho(\alpha_w+r))(\beta_{ww}(r-d)+\rho(\alpha_{ww}+r))}{\mu} \times$$

$$\frac{1}{\left(\begin{gathered} d^2\left(f_w h(\mu+\sigma_{ww})(\beta_{ww}(1-p)\rho+\beta_w\beta_{ww}(1-pq)+\beta_w p(1-q)\rho)-\right. \\ \left.\beta_w(\mu+\sigma_w)(\beta_{ww}(\mu+\sigma_{ww})-f_{ww}hpq(\beta_{ww}+\rho))\right)+ \\ d\left(f_w h(\mu+\sigma_{ww})(-\alpha_{ww}(1-p)\rho(\beta_w+\rho)\right. \\ -\alpha_w p(1-q)\rho(\beta_{ww}+\rho)-2\beta_w\beta_{ww}r(1-pq)+ \\ \beta_w\rho r(p(2q-1)-1)+\rho r(\beta_{ww}(pq+p-2)+\rho(pq-1)))+ \\ (\mu+\sigma_w)((\mu+\sigma_{ww})(\beta_{ww}\rho(\alpha_w+r)+\beta_w\rho(\alpha_{ww}+r)+ \\ \left.2\beta_w\beta_{ww}r)-f_{ww}hpq(\beta_{ww}+\rho)(\rho(\alpha_w+r)+2\beta_w r))\right)+ \\ f_w hr(\mu+\sigma_{ww})(\alpha_{ww}(1-p)\rho(\beta_w+\rho)+\alpha_w p(1-q)\rho(\beta_{ww}+\rho)+r(\beta_w+\rho)(\beta_{ww}+\rho)(1-pq))- \\ (\mu+\sigma_w)(\alpha_w\rho+r(\beta_w+\rho))(\alpha_{ww}\rho(\mu+\sigma_{ww})+r(\beta_{ww}+\rho)(-f_{ww}hpq+\mu+\sigma_{ww})) \end{gathered}\right)} \tag{SI.1}$$

and the reproduction rates $f_w$ and $f_{ww}$ satisfies either of the following conditions

$$f_{ww} \geq \frac{(\mu+\sigma_{ww})(-\alpha_{ww}\rho+\beta_{ww}d-r(\beta_{ww}+\rho))}{hpq(\beta_{ww}+\rho)(d-r)} \tag{SI.2}$$

or

$$f_{ww} < \frac{(\mu+\sigma_{ww})(-\alpha_{ww}\rho+\beta_{ww}d-r(\beta_{ww}+\rho))}{hpq(\beta_{ww}+\rho)(d-r)}$$

$$f_w > \frac{(\mu+\sigma_w)(-\alpha_w\rho+\beta_w d-r(\beta_w+\rho))}{h(d-r)(\mu+\sigma_{ww})} \times \tag{SI.3}$$

$$\frac{(r-d)(\beta_{ww}(\mu+\sigma_{ww})-f_{ww}hpq(\beta_{ww}+\rho))+\rho(\mu+\sigma_{ww})(\alpha_{ww}+r)}{\left(\begin{gathered} d(-\beta_{ww}(1-p)\rho+\beta_w\beta_{ww}(-(1-pq))-\beta_w p(1-q)\rho)+ \\ \alpha_{ww}(1-p)\rho(\beta_w+\rho)+\alpha_w p(1-q)\rho(\beta_{ww}+\rho)+r(\beta_w+\rho)(\beta_{ww}+\rho)(1-pq) \end{gathered}\right)} \tag{SI.4}$$

### SI 4. Equilibrium stability - Non-linear birth function for intermediate hosts

The disease-free equilibrium of the system of equations (1), (2), (3) given in the main text is

$$I_s^* = \frac{\mu}{c\rho} \tag{SI.5}$$

$$D_s^* = \frac{c\rho(r-d) - k\mu r}{c\rho^2} \tag{SI.6}$$

$D_s^*$ is positive if $X = c\rho(r-d) - k\mu r$ is positive.

The eigenvalues of the Jacobian matrix established at the disease-free equilibrium are roots of the following polynomial:

$$A_4\lambda^4 + A_3\lambda^3 + A_2\lambda^2 + A_1\lambda + A_0 \tag{SI.7}$$

The disease-free equilibrium is stable if the above polynomial has all negative real roots. Using the Descartes rule, the polynomial has all negative real roots if all the coefficients are positive.

We know that $A_4 = 1$ is always positive.

$$A_3 = \rho^7 \left( X(\beta_w + \beta_{ww} + 2\rho) + c\rho^2(2\alpha + \delta + \mu + \sigma + 2d) + \gamma\mu\rho \right) \tag{SI.8}$$

is always positive as all the elements of $A_3$ are positive.

$$\begin{aligned} A_2 =& \rho^{14} \left( c\rho^3(c\rho(\alpha^2 + 2\alpha(\delta + \mu + \sigma) + \delta(\mu + \sigma)) + (2\alpha + d + \mu + \sigma)(cd\rho + \gamma\mu) + cd\rho(2\delta + \mu + \sigma) + \gamma d\mu) \right. \\ & \left. + X(\beta_w + \beta_{ww} + 2\rho)(c\rho^2(\alpha + d + \delta + \mu + \sigma) + \gamma\mu\rho) + X^2(\beta_w + \rho)(\beta_{ww} + \rho) \right) \end{aligned} \tag{SI.9}$$

is always positive because all elements of $A_2$ are positive.

$$A_1 = \rho^{22}\left(c^2\rho^2 A_{10} + A_{11} - c\gamma X A_{12} f_w h\mu\rho^2\right) \tag{SI.10}$$

is positive if reproduction in single infection $f_w$, the probability to sucessfully established in the definitive host $h$, and cooperation in reproduction $\epsilon$ are small enough because

$$A_{10} = \alpha\rho^2(\alpha\gamma\mu + \alpha c\rho(\delta+\mu+\sigma) + 2(\mu+\sigma)(c\delta\rho+\gamma\mu)) \tag{SI.11}$$

is always positive, and

$$A_{11} = c\rho^2(2cd\rho\rho + X(\beta_w + \beta_{ww} + 2\rho))(\alpha\gamma\mu + \alpha c\rho(\delta+\mu+\sigma) + (\mu+\sigma)(c\delta\rho+\gamma\mu)) + \tag{SI.12}$$

$$cd\rho^2(c\rho(\delta+\mu+\sigma)+\gamma\mu)(cd\rho^2 + X(\beta_w+\beta_{ww}+2\rho)) + \tag{SI.13}$$

$$X^2(\beta_w+\rho)(\beta_{ww}+\rho)(c\rho(\delta+\mu+\sigma)+\gamma\mu) \tag{SI.14}$$

is always positive.

$$A_{12} = \beta_w(1-p) + p(\beta_{ww} + q(\epsilon-1)(\beta_{ww}+\rho)) + \rho \tag{SI.15}$$

is always positive because $0 \le p \le 1$ and $0 \le q \le 1$. If $\epsilon > 1$, then $A_{12}$ is always positive. The smaller the value of $\epsilon$, the more likely $A_{12}$ is negative. However, even when $\epsilon = 0$, $A_{12} = \beta_w(1-p) + \beta_{ww}p(1-q) + \rho(1-pq)$ is always positive. Because $A_{12}$ is positive, if $f_w$, $h$ and $\epsilon$ are sufficiently large, $A_1$ can be negative and the polynomial will not have all negative eigenvalues.

Finally, we have

$$A_0 = c^9\rho^{31}\left(-f_w h\gamma\mu A_{00} + A_{01}\right) \tag{SI.16}$$

where

$$A_{00} = X(c\rho^2(\alpha+d)A_{12} + X(\beta_w+\rho)(\beta_{ww}+\rho)(pq(\epsilon-1)+1)) \tag{SI.17}$$

is always positive and

$$A_{01} = (\mu+\sigma)(c\delta\rho+\gamma\mu)(c^2\rho^4(\alpha+d)^2 + cX\rho^2(\beta_w+\rho)(\alpha+d) + cX\rho^2(\beta_{ww}+\rho)(\alpha+d) + X^2(\beta_w+\rho)(\beta_{ww}+\rho)) \tag{SI.18}$$

is always positive. Therefore, if $f_w$ , $h$, and $\epsilon$ are sufficiently large, $A_0$ could be negative, leading to non-negative eigenvalues of the polynomial.

For all the above arguments, the disease-free equilibrium is positive and stable if $f_w$, $h$, and $\epsilon$ are sufficiently small, even though we cannot deduce the explicit expression for the condition.

### SI 5. Non-linear birth function for intermediate hosts - invasion condition

The condition for parasite invasion is $R_0 > 1$, which is satisfied when

$$f_w > \frac{\left(\begin{array}{c}(\mu+\sigma)(c\delta\rho+\gamma\mu)(k\mu r(\beta_w+\rho)-c\rho(\beta_w(-d)+\rho(\alpha+r)+\beta_w r))(k\mu r(\beta_{ww}+\rho)- \\ c\rho(\beta_{ww}(-d)+\rho(\alpha+r)+\beta_{ww}r))\end{array}\right)}{\left(\begin{array}{c}\gamma h\mu(c\rho(d-r)+k\mu r)(k\mu r(\beta_w+\rho)(\beta_{ww}+\rho)(pq(\epsilon-1)+1)- \\ c\rho(\beta_{ww}d(p-1)\rho-\beta_w d(\beta_{ww}+\beta_{ww}pq(\epsilon-1)+ \\ p\rho(q(\epsilon-1)+1))+\alpha\rho(\beta_w+\beta_w(-p)+ \\ p(\beta_{ww}+\beta_{ww}q(\epsilon-1)+q\rho(\epsilon-1))+\rho)+r(\beta_w+\rho)(\beta_{ww}+\rho)(pq(\epsilon-1)+1))\end{array}\right)} \tag{SI.19}$$

### SI 6. Effect of manipulation on the reproduction ratio

The effect of manipulation in single infection on the reproduction ratio $R_0$ is

$$\frac{dR_0}{d\beta_w} = \frac{\gamma D_s^* I_s^* f_w h(1-p)(d+\alpha_w)}{(\mu+\sigma_w)(\delta+\gamma I_s^*)(d+\alpha_w+D_{total}(\beta_w+\rho))^2} \tag{SI.20}$$

Both numerator and denominator are positive, therefore $R_0$ is an increasing function with respect to $\beta_w$.

The effect of manipulation in double infection on the reproduction ratio $R_0$ is

$$\frac{dR_0}{d\beta_{ww}} = \frac{\gamma D_s^* I_s^* f_w h p (d + \alpha_{ww})(\mu + (1-q)\sigma_{ww} + \mu q(\epsilon - 1) + q\sigma_w \epsilon)}{(\mu + \sigma_w)(\mu + \sigma_{ww})(\delta + \gamma I_s^*)(d + \alpha_{ww} + D_{total}(\beta_{ww} + \rho))^2} \tag{SI.21}$$

Both numerator and denominator are positive, therefore $R_0$ is an increasing function with respect to $\beta_{ww}$.

### Supplementary Figure

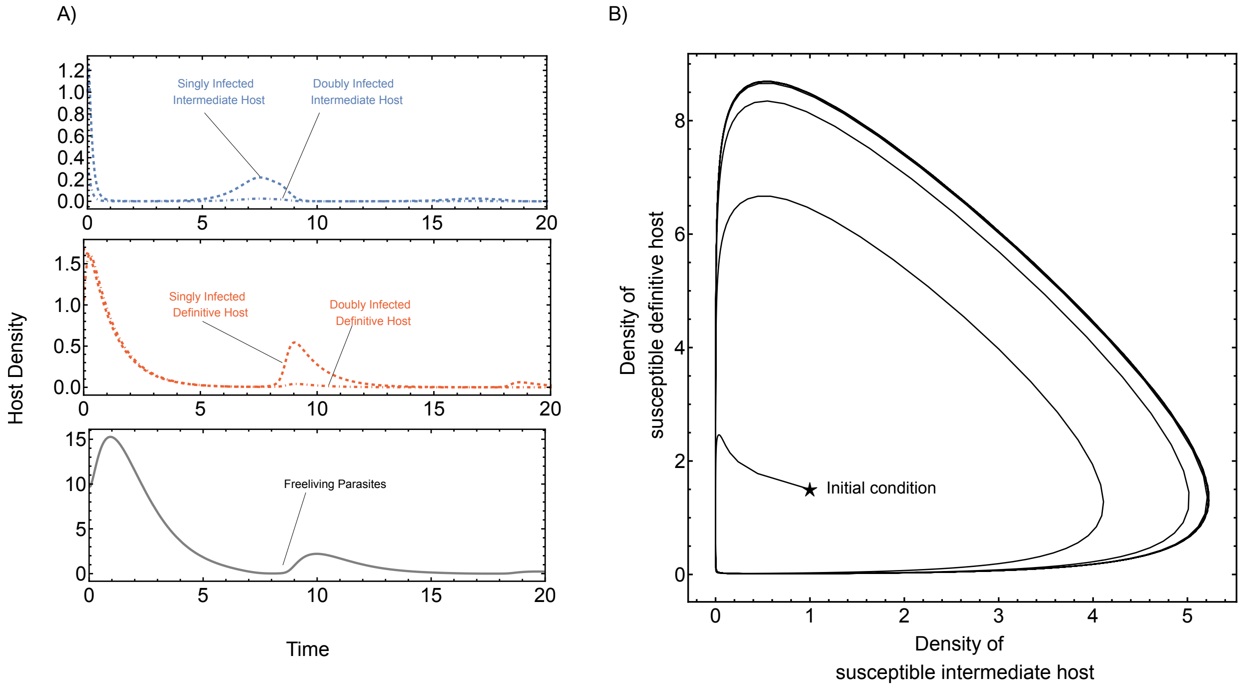


Figure SI.1: Disease-free equilibrium using a linear birth function, where parasite goes extinct (left panel), and susceptible hosts demonstrate cyclic dynamics (right panel). Solid grey lines indicate the density of free-living parasites, blue lines indicate infected intermediate hosts and red lines indicate infected definitive hosts. Dashed lines indicate singly infected hosts while dot-dashed lines indicate doubly infected hosts. Parameter values $\rho = 1.2$, $d = 0.9$, $r = 2.5$, $\gamma = 2.9$, $\alpha_w = \alpha_{ww} = 0$, $\beta_w = 1.5$, $\beta_{ww} = 1.5$, $p = 0.1$, $c = 1.4$, $\mu = 0.9$, $\sigma_w = \sigma_{ww} = 0$, $q = 0.01$, $f_w = 6.5$, $f_{ww} = 7.5$, $\delta = 0.9$,$h_1 = h_2 = 0.8$, $R_0 = 4.997$

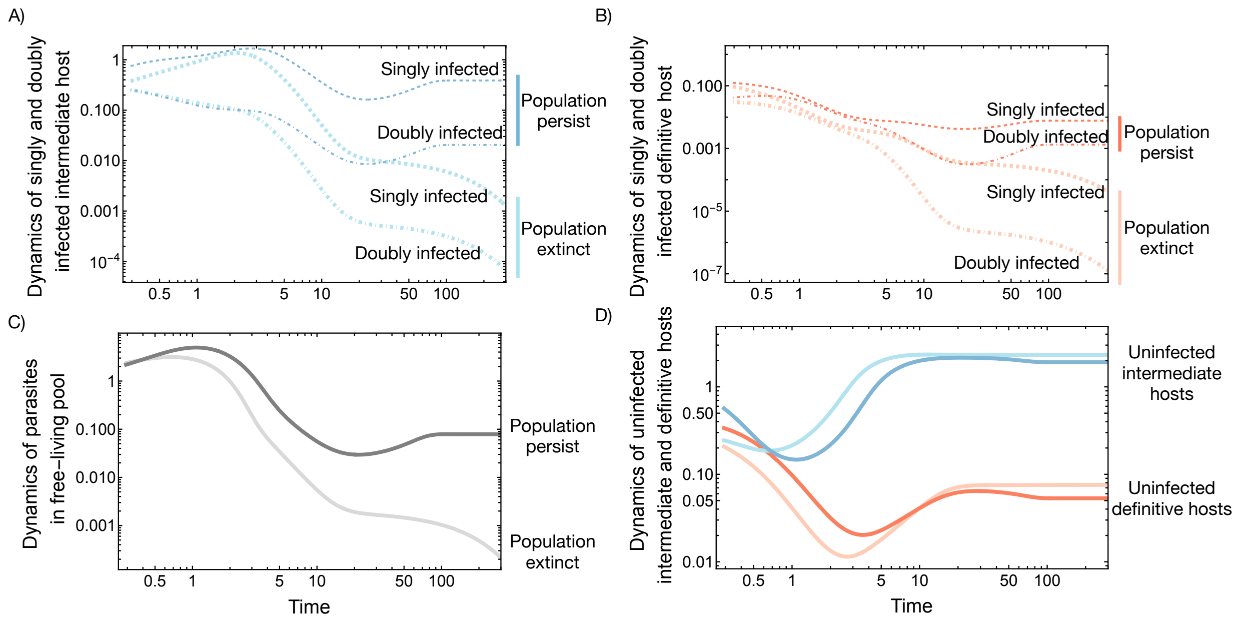


Figure SI.2: **Population dynamics of hosts and parasites.** A-C) Parasite populations go extinct when initial populations are insufficient and persist when initial populations are sufficient. D) Populations of unifected intermediate and definitive hosts always persist.

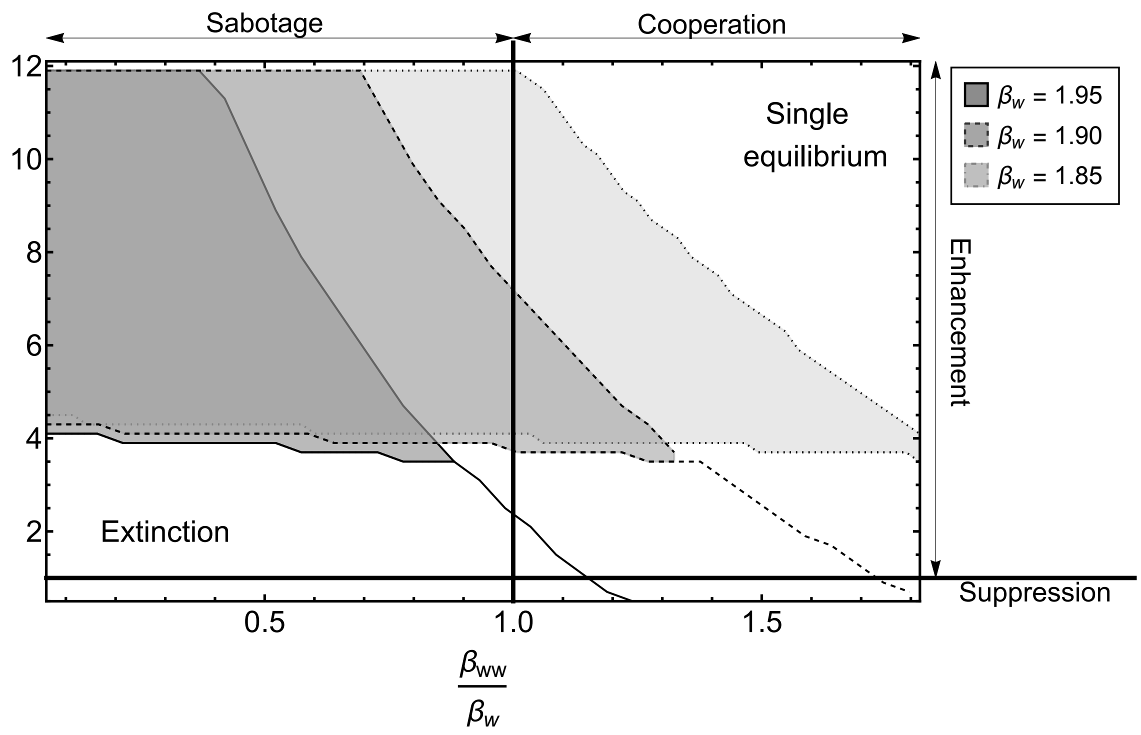


Figure SI.3: **Effect of manipulation and reproduction on bistability.** Changes in the bistability area (shaded areas) concerning different manipulation rates in single infection (different boundary styles). Manipulation and reproduction levels are equal between single and double infection on the vertical and horizontal lines. Common parameter: $\rho = 1.2$, $d = 0.9$, $r = 2.5$, $\gamma = 2.9$, $\alpha_w = 0$, $\alpha_{ww} = 0$, $p = 0.05$, $c = 1.4$, $\mu = 3.9$, $\sigma_w = 0$, $\sigma_{ww} = 0$, $q = 0.05$, $\delta = 0.9$, $k = 0.26$, $f_w = 35, h = 0.6$.
